# Enhancing CAR-T cell activity prediction via fine-tuning protein language models with generated CAR sequences

**DOI:** 10.1101/2025.03.27.645831

**Authors:** Kei Yoshida, Shoji Hisada, Ryoichi Takase, Atsushi Okuma, Yoshihito Ishida, Taketo Kawara, Takuya Miura-Yamashita, Daisuke Ito

## Abstract

Chimeric antigen receptor (CAR)-T cell therapy has shown remarkable success in treating hematological malignancies; however, several challenges remain, including limited efficacy against solid tumors, T cell exhaustion, and lack of T cell persistence, which have restricted its clinical efficacy across various indications. Sequence optimization of CAR constructs offers a promising strategy to enhance therapeutic efficacy of CAR-T cells. Recent advances in machine learning, especially protein language models (PLMs), enable prediction of mutational effects based on sequence representations. Nevertheless, applying PLMs to CARs is challenging due to the artificial nature of CARs and the absence of comprehensive CAR sequence databases. In this study, we developed a computational framework to predict CAR-T cell activity by fine-tuning ESM-2 with the CAR sequences generated using sequence augmentation. These CAR sequences were constructed by recombining homologous domains of CARs, enabling task-specific adaptation of the model. To evaluate prediction performance, we experimentally assessed the cytotoxicity of CAR-T cells expressing mutated CAR variants and compared these results with model predictions. Our results demonstrated that fine-tuned ESM-2 significantly improves prediction performance of CAR-T cell activity. Furthermore, we showed that training parameters—such as sequence diversity, number of training steps, and model size—substantially influence prediction performance. This work highlights the potential of combining sequence augmentation with fine-tuning PLMs to advance data-driven CAR-T cell design.

## Introduction

Chimeric antigen receptor (CAR)-T cell therapy holds great promise in revolutionizing cancer treatment. CARs are synthetic proteins that are combined distinct functional protein domains to recognize and eliminate target cells [1]. To date, seven CAR-T cell therapies have received FDA approval for hematological malignancies [2, 3]. However, CAR-T cell therapy has shown limited efficacy against solid tumors [4, 5], in part due to reduced persistence and attenuated anti-tumor effects [6, 7]. Addressing these challenges is essential for enhancing the therapeutic efficacy of CAR-T cells and expanding their application in cancer treatment.

Sequence optimization of CAR constructs represents a fundamental approach for improving the therapeutic performance of CAR-T cells [8]. One approach involves a introduction of point mutations into CAR constructs, which can significantly impact their function and efficacy. For instance, some mutations in both the single chain fragment variable (scFv) domain, which is responsible for modulating antigen binding affinity, and the intracellular domains, which control CAR-T cell activation and cytotoxicity, can enhance anti-tumor activity [9-12]. Therefore, designing CAR sequences with desired therapeutic outcomes requires a systematic introduction of diverse mutations across CAR domains and a comprehensive evaluation of their effects.

Machine learning (ML) techniques are increasingly recognized as powerful tools for predicting the mutational effects in various proteins [13-15]. Most ML models learn the relationship between numerical representations derived from amino acid sequences and their functional characteristics [16]. In particular, protein language models (PLMs) have attracted significant attention for their ability to extract meaningful representations from amino acid sequences [17].

PLMs are ML models, often based on deep learning architectures, that employ techniques similar to those utilized in natural language processing [18-21]. Trained on millions of amino acid sequences, PLMs can capture evolutionary patterns and structural relationships inherent to natural proteins. To obtain meaningful sequence representations for specific proteins, fine-tuning the parameters in PLMs is a commonly beneficial approach. Fine-tuning involves adapting pre-trained PLMs to particular tasks by further training it on smaller, task-specific datasets [22-24]. For example, evolutionary fine-tuning (evotuning) is an unsupervised learning approach that utilizes homologous sequences of a target protein [18]. Fine-tuned models obtained via evotuning can capture evolutionary relationships and functionally relevant patterns in amino acid sequences, thereby enhancing their utility in downstream prediction tasks [25-27]. In another study, Vincoff *et al*. demonstrated that fine-tuning a PLM using sequences of fusion oncoprotein—a class of chimeric proteins—collected from databases led to superior performance compared to baseline embeddings in various fusion-specific tasks [28]. However, it is challenging to directly utilize the evotuning approach for analysis of CAR sequences because CARs are synthetic proteins. Additionally, the absence of focused CAR sequence databases and the limited availability of reported CAR sequences further complicate fine-tuning PLMs for CARs. These factors hinder accurate prediction of CAR-T cell activity using PLMs.

In this study, we newly generated unlabeled CAR sequences through sequence augmentation and developed prediction models for CAR-T cell activity by fine-tuning PLMs with the generated sequences. To generate a large and diverse set of unlabeled CAR sequences, homologous sequences from individual domains of a target CAR region (Fig 1A) —specifically CD28 and CD3ζ—were randomly combined. The generated sequences were then used to fine-tune ESM2, a PLM developed by Meta AI, available in various model sizes [29]. To validate whether our methodology is effective for predicting CAR-T cell activity, numerous CAR mutants were constructed by introducing point mutations into a wild-type sequence (S1 Table), and the cytotoxicity of CAR-T cells expressing these mutants was experimentally assessed. Subsequently, the cytotoxicity was predicted based on CAR sequences using the fine-tuned ESM2 (Fig 1B). Our results demonstrated that the fine-tuned ESM2 significantly improved the prediction performance of CAR-T cell activity. Furthermore, we showed that optimizing the fine-tuning conditions—such as sequence diversity, the number of training steps, and the choice of ESM2 model size—can maximize the fine-tuning efficacy and enhance the prediction performance.

**Fig 1.**
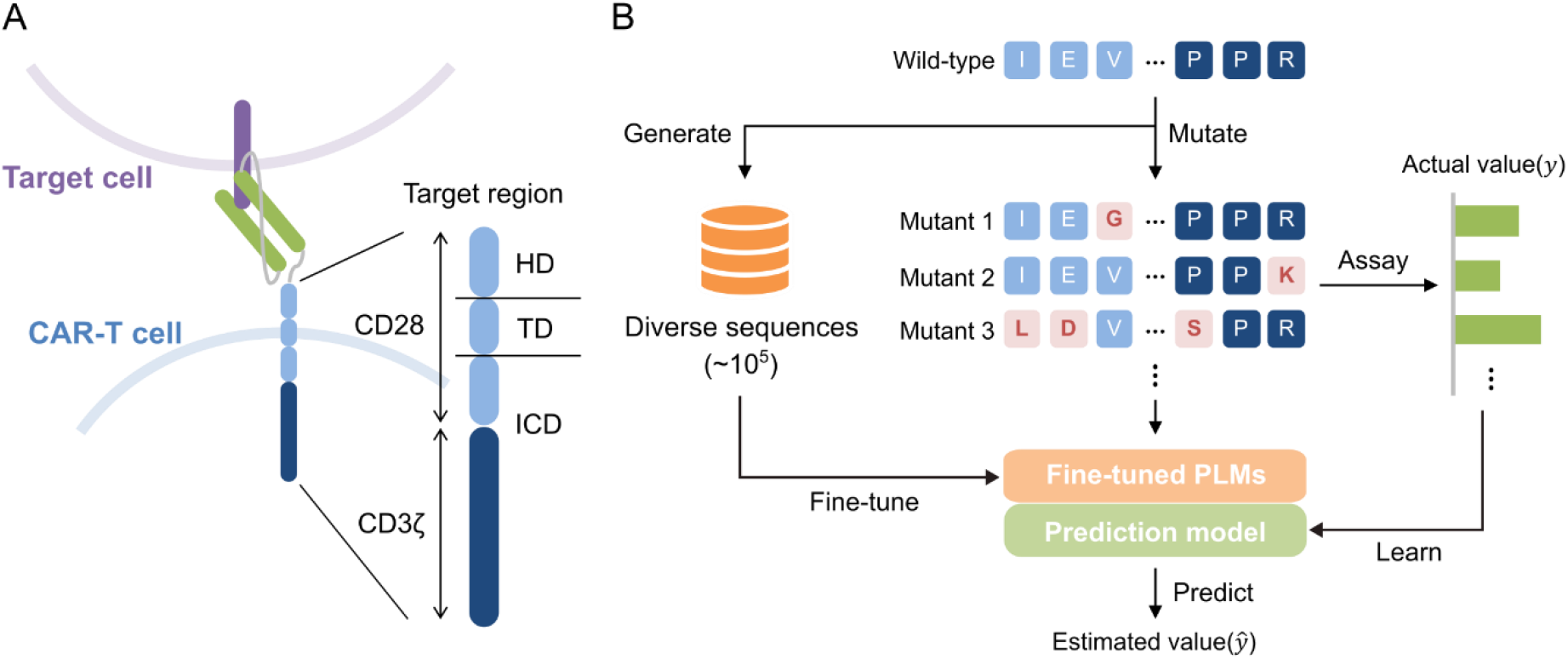
Schematic outline of our methodology. (A) Illustration of a target CAR region in this study. We focused on the hinge domain (HD), transmembrane domain (TD), and intracellular domain (ICD) which are derived from CD28 and CD3ζ. (B) Scheme for predicting CAR-T cell activity. The CAR mutants were obtained by introducing point mutations into the target CAR region, and the cytotoxicity of CAR-T cells was experimentally evaluated. In addition, unlabeled CAR sequences were constructed using the wild-type sequence as a template. These generated sequences were used to fine-tune the PLMs, and subsequently the fine-tuned PLMs were used to train the prediction model.

## Results

### Sequence Augmentation for Fine-Tuning PLMs: High-diversity and Low-diversity Sequence Groups

To fine-tune pre-trained PLMs for predicting CAR-T cell activity, a large number of CAR sequences are required. Therefore, we generated numerous unlabeled CAR sequences through sequence augmentation. In this study, we focused on a chimeric region composed of CD28 and CD3ζ within a target CAR sequence. In the sequence augmentation process, homologous sequences for each protein domain were collected using HMMER software [30] and combined randomly. Previous studies have shown that human 4-1BB and subunits of CD3 are functional in mouse T cells [31-33]. Additionally, CAR with CD3ζ replaced by subunits of CD3 also exhibited functionality [34]. Hence, we hypothesized that utilizing homologous sequences could be applicable in this context. Our methodology of sequence augmentation facilitated the creation of a diverse set of sequences based on different combination patterns. It is well known that the diversity of training data has impact on the prediction results, and that a high level of diversity is expected to contribute to improved performance [29]. Therefore, we established two sequence groups with distinct characteristics: a high-diversity group and a low-diversity group (details provided in Materials and Methods) (Table 1). To visualize the pattern of the generated sequences, they were first one-hot encoded and then subjected to dimensionality reduction using multi-dimensional scaling (MDS) (Fig 2A). The high-diversity group exhibited a distribution across the entire sequence space, whereas the low-diversity group clustered around the wild-type sequence. Additionally, the distribution of sequence lengths and the Levenshtein distances [35] from the wild-type sequence are shown in Fig 2B and 2C. The Levenshtein distance denotes the similarity between two sequences, defined as the minimum number of single-character edits (insertions, deletions, or substitutions) required to transform one sequence into the other. Compared to the low-diversity group, the high-diversity group exhibited a broader range of sequence lengths and lower similarities to the wild-type sequence. These results confirmed the successful establishment of two sequence groups characterized by distinct levels of diversity.

**Table 1.** Comparison of generated sequences with different diversity.

| Diversity | Number of sequences | Length |  |  | Levenshtein distance |  |  |
| --- | --- | --- | --- | --- | --- | --- | --- |
|  |  | Mean | Max | Min | Mean | Max | Min |
| High | 5500 | 211 | 237 | 170 | 87 | 153 | 0 |
| Low | 5250 | 218 | 226 | 180 | 45 | 91 | 0 |

**Fig 2.**
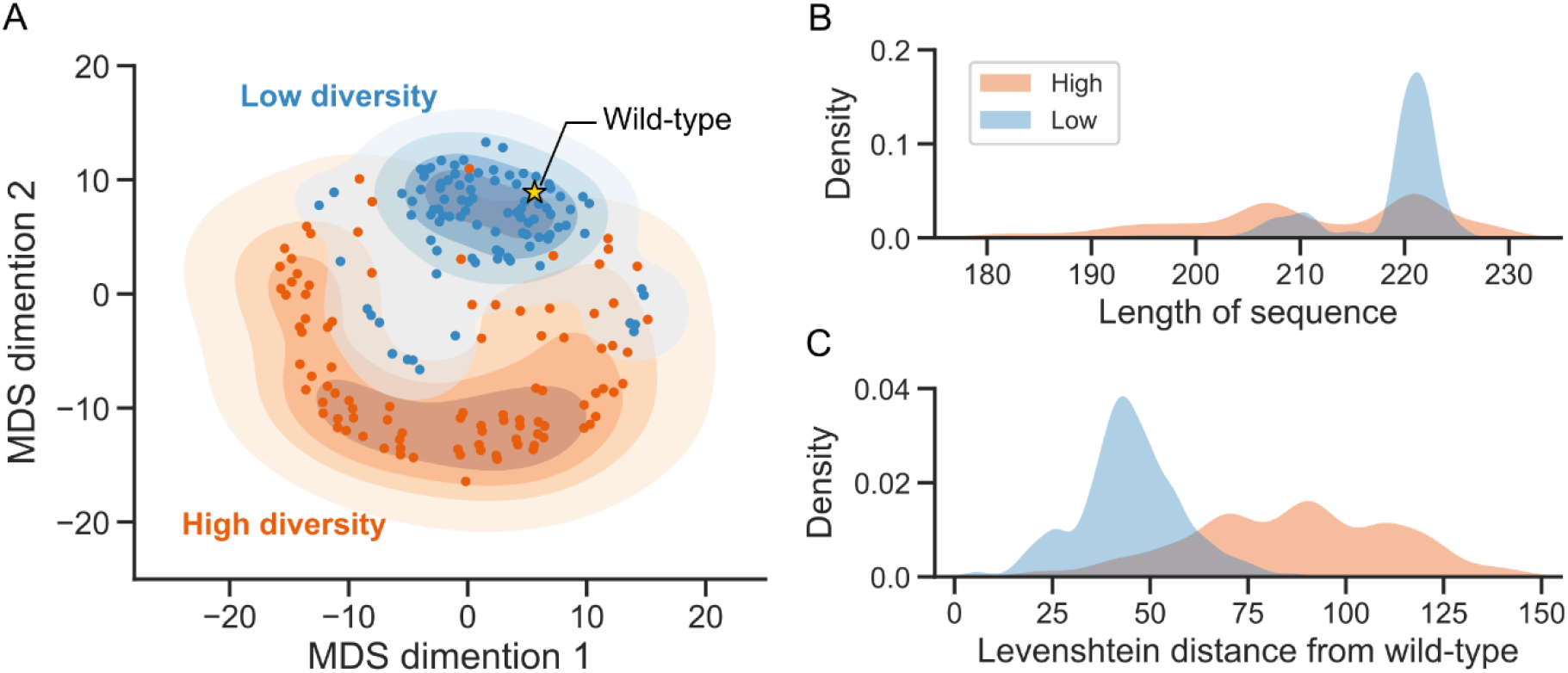
Analysis of generated CAR sequences. (A) MDS plot of generated sequences, comparing the high-diversity and low-diversity groups. The orange plots represent sequences belonging to the high-diversity group, while the blue plots represent sequences belonging to the low-diversity group. The filled contours represent the kernel density estimate (KDE). The yellow star indicates the wild-type (= original sequence before introducing mutations) sequence. (B) Distribution of the sequence lengths. (C) Levenshtein distances from the wild-type sequence.

### Data Acquisition of CAR-T Cell Activity Associated with Mutated CAR Sequences

To evaluate the impact of fine-tuning on the prediction of CAR-T cell activity, we obtained data on mutated CAR sequences and their cytotoxicity. Initially, mutated CAR sequences were acquired by introducing point mutations into CD28 and CD3ζ in a wild-type CAR sequence, resulting in a total of 382 CAR mutants. The number of mutations per sequence ranged from 1 to 10, with the majority containing 10 mutations (Fig 3A). Subsequently, the cytotoxicity of CAR-T cells expressing these CAR mutants was investigated. The cytotoxicity of 382 mutants is distributed over the entire range, with that of the wild-type around approximately 50% (Fig 3B). Thus, we have successfully acquired the CAR mutants and their cytotoxicity data, enabling the evaluation of fine-tuning effects.

**Fig 3.**
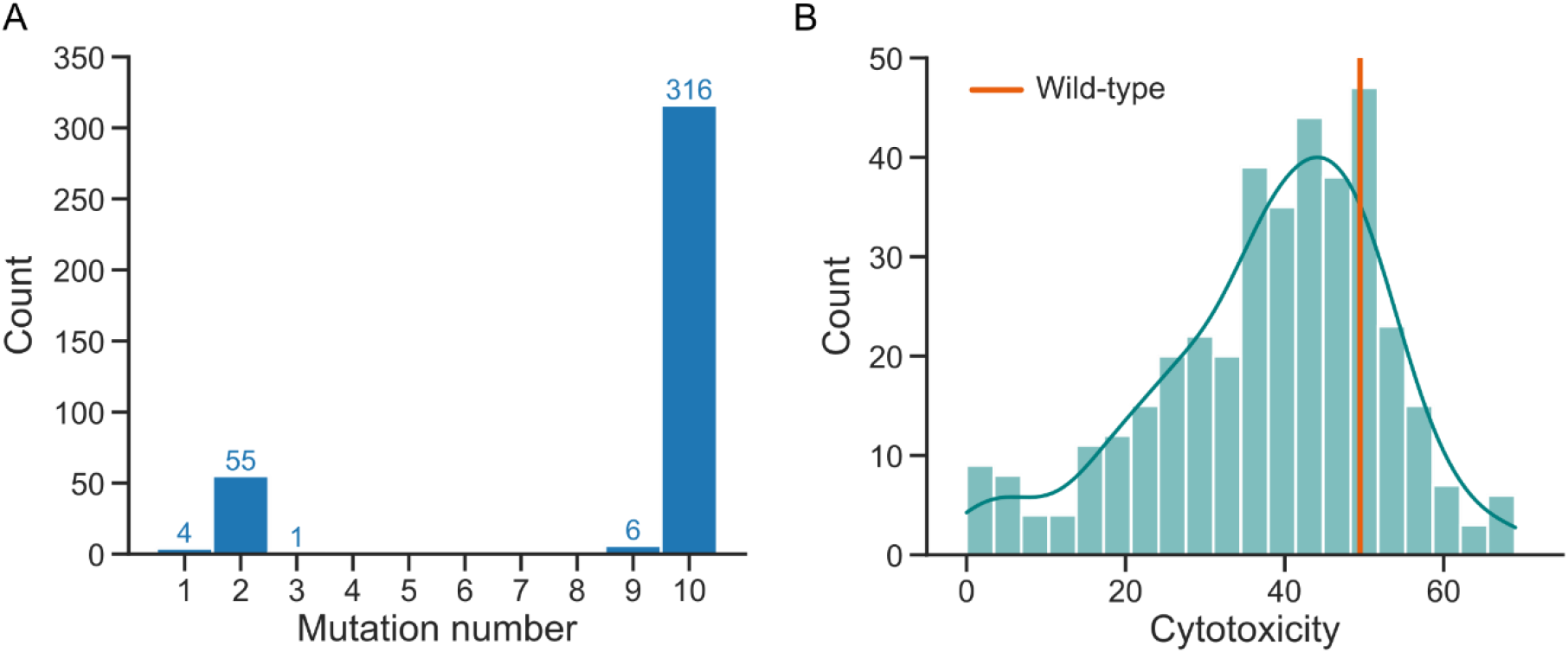
Visualization of CAR mutant dataset. (A) Distribution of the mutation number per mutated sequence. The numbers above each bar indicate the count, and there are no sequences with mutation number 4 to 8. (B) Distribution of cytotoxicity in a total of 382 CAR mutants. The vertical orange line indicates cytotoxicity of wild-type sequence.

### Fine-Tuning ESM2 and Predicting Cytotoxicity of CAR-T Cells Using Generated CAR Sequences

To determine the optimal model size of ESM2 for fine-tuning, we evaluated four pre-trained ESM2 models (S2 Table) and compared their prediction performance for the cytotoxicity of the CAR mutants. For prediction, we chose a Ridge regression, which is a linear regression model that incorporates regularization to reduce overfitting, thereby enhancing its performance on high-dimensional data such as amino acid sequence representation [26]. Among the evaluated models, ESM2-35M exhibited the most superior prediction performance, while the other models consistently showed lower prediction performance (S1 Fig). Consequently, we concluded that ESM2-35M was the optimal model to develop higher performing prediction model through fine-tuning^1^.

Next, we performed cytotoxicity prediction using the fine-tuned ESM2-35M model. Fine-tuning was executed for up to 50 epochs on the sequences from both the high-diversity and low-diversity groups. Loss and perplexity against training and validation data gradually decreased towards 50 epochs in both groups (S2 Fig). The relationship between the number of epochs and Spearman’s rank correlation coefficient was illustrated in Fig 4A, B. In both groups, prediction performance peaked within the initial training steps, with the high-diversity group peaking at 10 epochs and the low-diversity group at 5 epochs. Each prediction model was defined as ESM2-35M_H/e10_ and ESM2-35M_L/e5_, respectively. Moreover, ESM2-35M_H/e10_ led to a significant improvement in prediction performance, increasing by 20.2% (from 0.318 to 0.458) (Fig 4C, Table 2), with performance improvement observed across all splits in the cross-validation (S3 Fig).

**Table 2.**
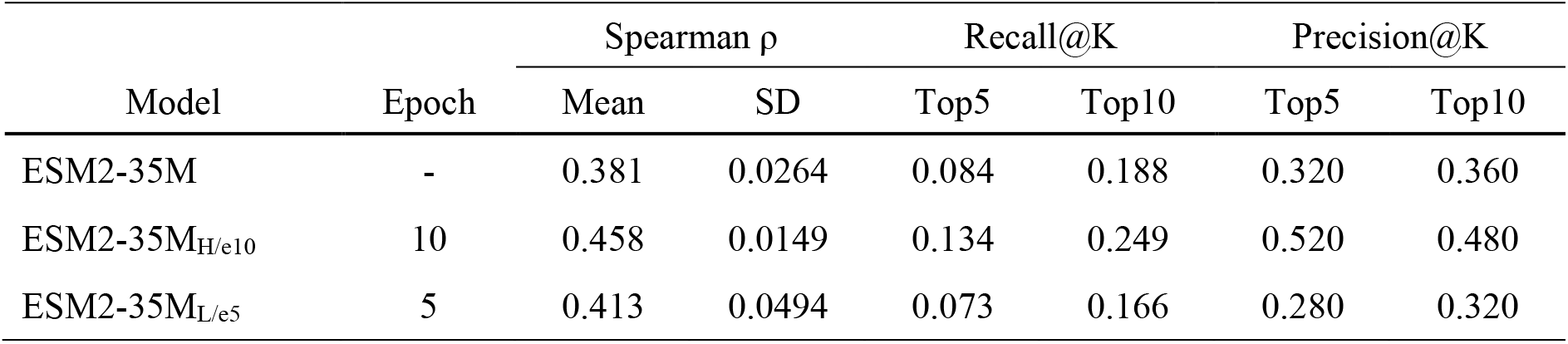
Summary of prediction performance using pre-trained and fine-tuned ESM2-35M.

| Model | Epoch | Spearman $\rho$ | | Recall@K | | Precision@K | |
| --- | --- | --- | --- | --- | --- | --- | --- |
|  |  | Mean | SD | Top5 | Top10 | Top5 | Top10 |
| ESM2-35M | - | 0.381 | 0.0264 | 0.084 | 0.188 | 0.320 | 0.360 |
| ESM2-35M<sub>H/e10</sub> | 10 | 0.458 | 0.0149 | 0.134 | 0.249 | 0.520 | 0.480 |
| ESM2-35M<sub>L/e5</sub> | 5 | 0.413 | 0.0494 | 0.073 | 0.166 | 0.280 | 0.320 |

**Fig 4.**
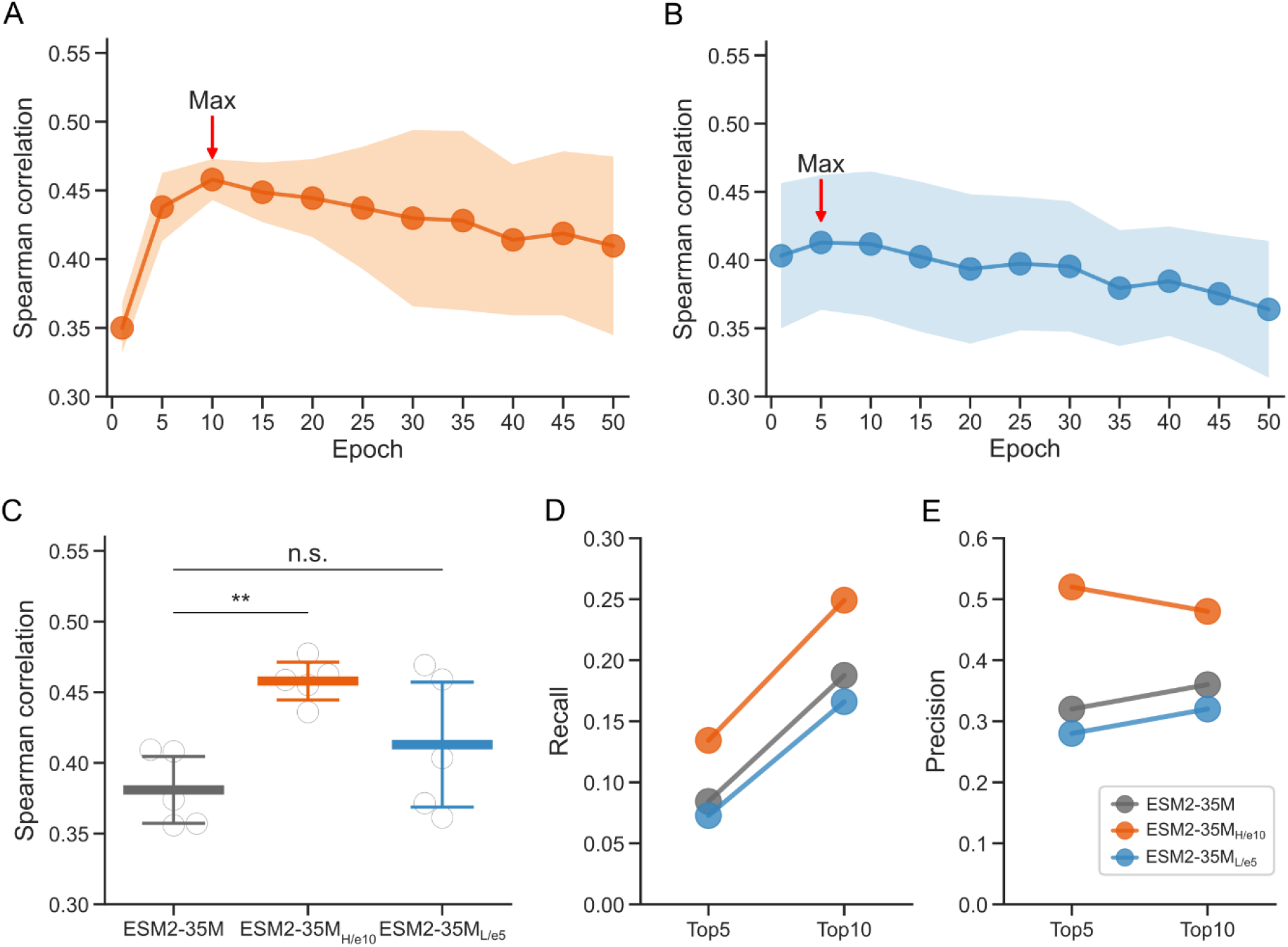
Prediction performance of fine-tuned ESM2-35M. (A, B) The relationship between the number of epochs and Spearman’s rank correlation coefficient. The plots indicate the average value for five models generated by cross-validation, and the filled regions show the standard deviation. The red arrows indicate the maximum value within the range of 5 to 50 epochs. (C) Comparison of Spearman’s rank correlation coefficient between pre-trained and fine-tuned ESM2-35M. The results of the fine-tuned ESM2-35M were taken at the maximum point in (A). White circles represent individual values for each model of cross-validation. The thick middle lines represent the mean values, and the thin upper and lower lines indicate the mean ± SD. Multiple comparisons were performed using Dunnett’s test, **p<0.01, n.s. (not significant). (D, E) Recall@K and Precision@K for K=5 and K=10.

In addition, binary evaluation was performed to validate the capability of ESM2-35M_H/e10_ in predicting high-cytotoxicity CARs. Accurate prediction of top-ranked data is crucial for designing proteins with enhanced activity [30]. Specifically, the top 25% of cytotoxicity in the validation data were classified as the high-cytotoxicity CARs, and both Recall at K (Recall@K) and Precision at K (Precision@K) scores were calculated. These metrics are commonly used for evaluation in ranking and retrieval tasks. Recall@K indicates how effectively the model retrieves all high-cytotoxicity CARs, while Precision@K signifies how many of the top K predictions are high-cytotoxicity CARs. The evaluation results for K=5 and K=10 revealed that both Recall@K and Precision@K improved with ESM2-35M_H/e10_ (Fig 4D, E). These findings demonstrated that fine-tuning with the sequences from the high-diversity group enhanced the prediction performance of CAR-T cell cytotoxicity.

## Discussion

PLMs are valuable tools for extracting meaningful representations from amino acid sequences. Fine-tuning pre-trained PLMs on task-specific datasets can enhance prediction performance in various bioinformatics applications. Nevertheless, fine-tuning PLMs for predicting CAR-T cell activity poses notable challenges given the artificial nature of CARs and the lack of focused CAR sequence databases. In this study, we generated unlabeled CAR sequences and investigated the impact of fine-tuning ESM2 using these sequences for downstream prediction tasks. Evotuning has been shown to be an effective approach for fine-tuning PLMs, enhancing their performance through training on homologous sequences [18]. Our method of sequence augmentation, which involves combining homologous sequences, is inspired by this approach. While CARs are synthetic proteins, the homologs of the domains in CARs have shown CAR-T cell function [31, 34]. Therefore, we anticipated that our approach of utilizing evolutionary sequences would be effective.

In sequence augmentation, clustering was performed utilizing the characteristics of homologous sequences collected from database, and two types of sequences were created by randomly combining them from distinct clusters. This methodology has the capacity to generate a large and diverse set of sequences simply by adjusting the number of clusters and the manner in which they are combined. Consequently, our approach holds potential for broader applications in the analysis of other multi-domain proteins in the future research.

Among the pre-trained ESM2 models utilized in this study, ESM2-35M exhibited the highest prediction performance for our specific task (S1 Fig). The size of PLMs is directly linked to their performance in downstream tasks, with larger models typically offering superior predictive capabilities [38, 39]. Nonetheless, the advantage of scaling up model size depends on the task complexity, and smaller models may already provide sufficient performance for certain tasks [25]. In our study, ESM2-35M emerged as the most appropriate choice for predicting CAR-T cell activity. These results highlight that task-specific requirements and data considerations are critical in determining the optimal model size. In addition, ESM2-35M_H/e10_ exhibited a statistically significant improvement in prediction performance as a result of fine-tuning. In contrast, the other models not only exhibited lower prediction performance but also failed to show any improvement through fine-tuning (S5 Fig, S3 Table). These results imply the significance of selecting an appropriate model size for PLMs to effectively utilize the benefits of fine-tuning.

We showed that only the high-diversity group significantly enhanced the prediction performance for CAR-T cell cytotoxicity (Fig 4C). This result was consistent with a previous report hypothesizing that the high-diversity of the training data improves the model’s robustness [36]. Given the absence of CAR sequences in the pre-training data of ESM2, the high-diversity of sequences may have enabled the model to gain insight into a broader range of sequence-function relationships. These findings suggest that adjusting sequence diversity for fine-tuning according to each downstream task is essential to maximize prediction performance. Furthermore, in contrast to the prediction performance, both ESM-35M_H/e10_ and ESM-35M_L/e5_ showed similar log-likelihood values for CAR mutants (S4 Fig). A previous study has highlighted that pre-training performance does not consistently correlate with downstream task performance [37]. Hence, the diversity of the training data does not notably enhance the log-likelihood of CARs, but it plays a crucial role in predicting CAR-T cell activity.

During the fine-tuning process of ESM2-35M using the high-diversity group, prediction performance reached its peak at 10 epochs (Fig 4A). Beyond this point, performance gradually decreased, and variance increased with additional epochs, indicating overfitting in certain cross-validation splits during the later stages of fine-tuning. The limited availability of training data and the use of the mask replacement task may contribute to overfitting [40]. Therefore, excessive fine-tuning on small datasets could compromise generalization, whereas selecting an optimal number of training steps has the potential to enhance prediction performance while minimizing overfitting.

It has been reported that adjusting affinity through point mutations in the scFv can enhance therapeutic efficacy [9, 10]. In addition, PLMs for hit maturation of scFv are valuable for discovering candidate CARs [41]. Our approach remains applicable even as the number of target domains expands, suggesting the potential for future research to optimize the full-length CAR, including the scFv region. Futhermore, our study demonstrated that sequence augmentation with different diversity, coupled with fine-tuning the ESM2 model, can significantly improve the prediction performance for the cytotoxicity of CAR-T cells. In the future, combining our methodology with high-throughput screening of CAR mutants could facilitate the efficient design of novel CAR sequences, thereby enabling enhancement and selection of CAR-T cell candidates.

## Materials and Methods

### Sequence augmentation

We utilized the HMMER software (https://www.ebi.ac.uk/Tools/hmmer/) [30] to search for homologous sequences of CD28 (position: 115-220) and CD3ζ (position: 52-164) using the phmmer algorithm. The E-value threshold was set to 1e-4, and Uniprot was employed as the database. After removing duplicates, sequences that were longer than ×1.25 or shorter than ×0.75 the length of the wild-type were filtered out. Subsequently, the sequences were converted into numerical vectors using one-hot-encoding, and K-means clustering was performed for each domain. The number of clusters (K) was determined to ensure that at least 25% of the total sequences were included in clusters containing wild-type sequences. In this study, K was set to 4 for CD28 and 3 for CD3ζ. The sequences in high-diversity group were generated by randomly sampling and combining sequences from each cluster of CD28 and CD3ζ, whereas the sequences in low-diversity group were generated by sampling and combining sequences from clusters containing the wild-type sequences of CD28 and CD3ζ. In this study, the high-diversity group comprised 5,500 sequences, and the low-diversity group comprised 5,250 sequences. The scheme was illustrated in S6 Fig.

### CAR mutants

We used anti-CD19 FMC63-CAR, which includes FMC63 scFv, CD28, and CD3ζ domains, as the template sequence. Random point mutations, up to a total of 10, were introduced into the hinge, transmembrane, and intracellular domains of CD28 and CD3ζ. As a result, we acquired a total of 382 CAR mutants.

### Cell culture

Two donors of primary human CD8 T cells (STEMCELL technologies) were cultured with X-VIVO15 (LONZA) with 5% human serum (Sigma-Aldrich), 100 μM 2-Mercaptoethanol (FUJIFILM-Wako), 25 U/mL human IL-2 (Peprotech), 10 ng/mL human IL-7 (Miltenyi Biotec), and 10 ng/mL human IL-15 (Miltenyi Biotec).

### Transduction

eBlocks genes (Integrated DNA Technologies) encoding CAR mutant sequence were inserted into pHR-SFFV plasmid vectors by the Gibson Assembly method. Then, the E. coli which transformed plasmid vectors was cultured for vector expansion. Vector purification was performed with miniprep kit (Qiagen). HEK293FT cells (Thermo Fisher Scientific) were transfected with pHR-GOI vector and packaging plasmid mix (pMD2.G, pCMVR8.74, and pAdvantage) to produce replication incomplete lentivirus vector. The viral supernatant was harvested after 2 days. Human CD8 T cells were stimulated with Human T-activator CD3/CD28 Dynabeads (Thermo Fisher Scientific) at a 1:1 cell-to-bead ratio and culture for 24 hours and then mixed with the viral supernatant. After culturing for 2 days, the beads were removed from T cell culture with Magnum FLX magnet plate (ALPAQUA). Two days later, the cells were used for cytotoxic assay.

### Cytotoxicity assay

Cytotoxicity assays were performed using a bioluminescence-based method. In brief, Nalm-Luc cells were seeded in each well of a 384-well round bottom plate (Sumitomo Bakelite). Subsequently, CAR-T cells were added at an effector-to-target (E:T) ratio of 1:8. After co-culturing for 23 hours, luciferin reagent (OneGlo, Promega) was added at half the volume of the total culture medium, and luminescence was measured using a microplate reader (SpectraMax i3x, Molecular Devices). Target cell cytotoxicity was determined using the following formula:

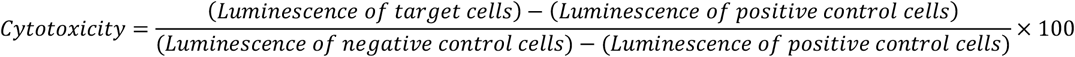

The cytotoxicity values for each mutant were calculated based on data from two donors, and the average of these values was calculated.

### Statistical analysis

Dunnett’s multiple comparison test was performed to assess the significance of the differences in the Spearman’s rank correlation coefficient in each split of the 5-fold cross-validation between pre-trained and fine-tuned ESM2. The Dunnett’s test was executed utilizing the Python package “scipy.stats.dunnett”.

### Multi-dimensional scaling analysis

Multi-dimensional scaling (MDS), a statistical technique used to reduce the dimensionality of data, was performed to visualize the generated sequences. Prior to the MDS analysis, 2% of the sequences from both the high-diversity and low-diversity groups were randomly sampled. These sequences were subsequently transformed into numerical vectors using one-hot encoding. The MDS analysis was conducted utilizing the Python package “sklearn.manifold.MDS”.

### Fine-tuning

We conducted full model fine-tuning using masked language modeling. The PyTorch framework [42] and the Hugging Face transformers library [43] were used to implement the code for loading and fine-tuning the ESM-2 model. Two datasets, high-diversity and low-diversity groups, were used for fine-tuning. Prior to fine-tuning, the dataset was split into 80% for training and 20% for validation. The AdamW optimizer was used with a learning rate of 5e-6 for 50 epochs, and the batch size was set to 128. Fine-tuning was performed on 8 NVIDIA V100 graphical processing units (GPUs). The fine-tuned models were saved every 5 epochs and used in the downstream task.

### Pseudo log-likelihood

Wang and Cho proposed the pseudo log-likelihood (PLL) for masked language models [44]. PLL was calculated for each position in the target sequence when masked, and the average logarithmic likelihood was then computed. The PLL was calculated using the following formula:

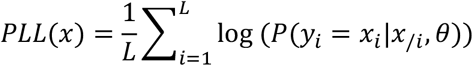

In this study, we calculated the PLLs of 382 CAR mutant sequences using pre-trained ESM2-35M and fine-tuned ESM2-35M with high-diversity and low-diversity groups, and then averaged the results.

### Sequence representation

A sequence containing L amino acids was input into the pre-trained PLMs, resulting in a numerical matrix. The numerical matrix obtained from each layer of PLMs was of size (L+2) × H. The addition of 2 accounts for the special tokens [CLS] and [EOS], which were added at the beginning and end of the sequence, respectively. H represents hidden size, which corresponds to the size of the individual embedding by each PLM (S2 Table). We extracted fixed-size vectors from the [CLS] token of the last layer to serve as sequence features.

### Prediction model

The prediction models were constructed using sequence representations to predict the cytotoxicity of CAR-T cells. In this study, we employed a Ridge regression model, which is a type of linear regression that includes a regularization term. To assess the performance of the prediction model, nested cross-validation was used. Briefly, prior to prediction, the dataset was divided into five splits, with four of the five used as training data and the remaining one used as validation data, resulting in the creation of five models (5-fold cross-validation in outer loop). The prediction scores, which represent the correlation between the predicted values and the actual values, were calculated for the five models. The hyperparameter of the Ridge model, regularization strength(α), was optimized over the range of 10^−6^ to 10^6^ using the training data divided into three splits (3-fold cross-validation in the inner loop), and the best model was used for the prediction of the validation data. We utilized the Python package “sklearn.linear_model.RidgeCV” with cv=3 for model implementation.

### Prediction score

To evaluate the prediction models, two types of prediction scores were used in this study. First, we calculated Spearman’s correlation coefficient between the estimated and actual values to assess the overall trend of the predictions. Second, we calculated Recall at K (Recall@K) and Precision at K (Precision@K) to evaluate whether high-cytotoxicity CAR mutants could be accurately predicted. Briefly, Recall@K evaluates how effectively the model retrieves all high-cytotoxicity CARs, whereas Precision@K evaluates how many of the top K predictions are high-cytotoxicity CARs. In this study, we labeled the top 25% of cytotoxicity values in the validation data as the high-cytotoxicity CARs. Both Recall@K and Precision@K were calculated for K=5 and K=10 using the following formulas:

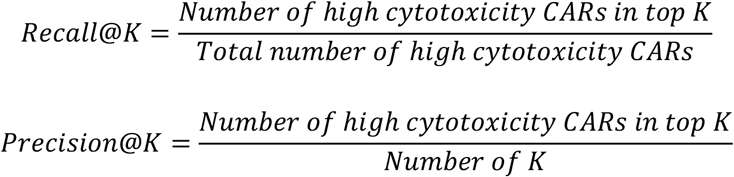

## Acknowledgements

The authors would like to thank Satoru Mimura for technical assistance, and Kenichi Hironaka, Tomohiko Okuda, Hiroya Ijima, Yoshihiro Osakabe, and Akinori Asahara for helpful discussion. Many thanks go to my supervisors, Shizu Takeda and Hiroko Hanzawa, for advice, encouragement, and support.

## Author contributions

**Kei Yoshida:** Writing – original draft, Conceptualization, Data curation, Formal analysis, Methodology, Software, Validation, Visualization, **Shoji Hisada:** Writing – review & editing, Conceptualization, Methodology, **Ryoichi Takase:** Writing – review & editing, Software, Validation, **Atsushi Okuma:** Writing – original draft, Writing – review & editing, Investigation, Resources, **Yoshihito Ishida:** Writing – review & editing, Investigation, Resources, **Taketo Kawara:** Writing – review & editing, Investigation, Resources, **Takuya Miura-Yamashita:** Writing – review & editing, Investigation, Resources, **Daisuke Ito:** Writing – review & editing, Investigation, Resources.

## Supporting information

**S1 Fig.**
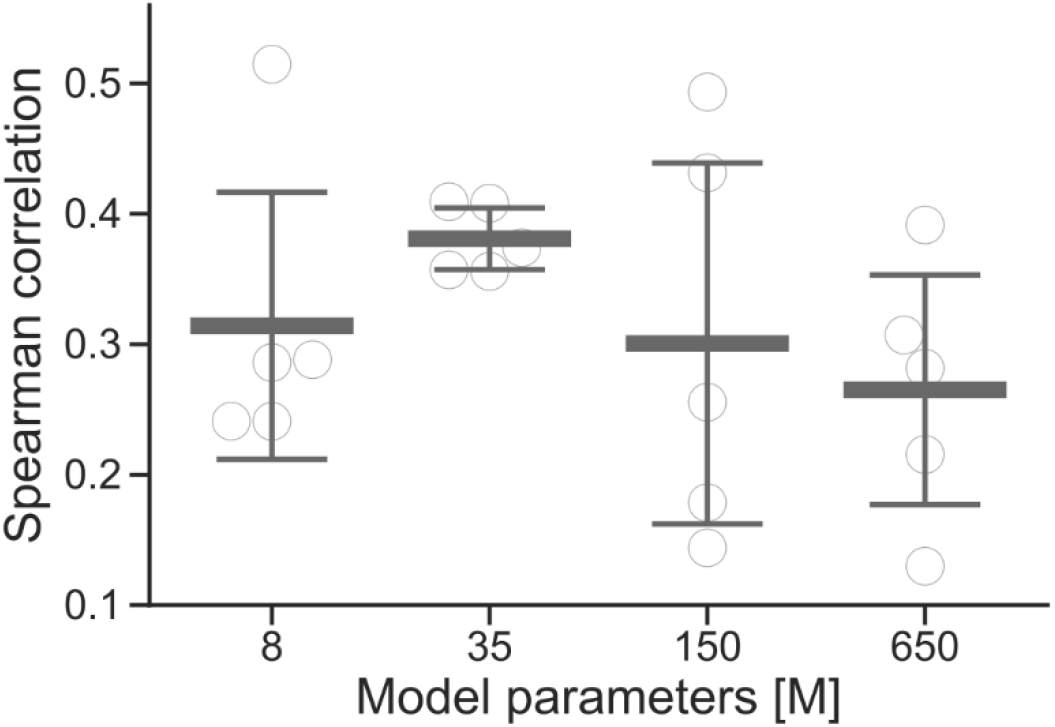
Comparison of Spearman’s rank correlation coefficient between pre-trained ESM2. White circles represent individual values for each model of cross-validation. The thick middle lines represent the mean values, and the thin upper and lower lines indicate the mean ± SD.

**S2 Fig.**
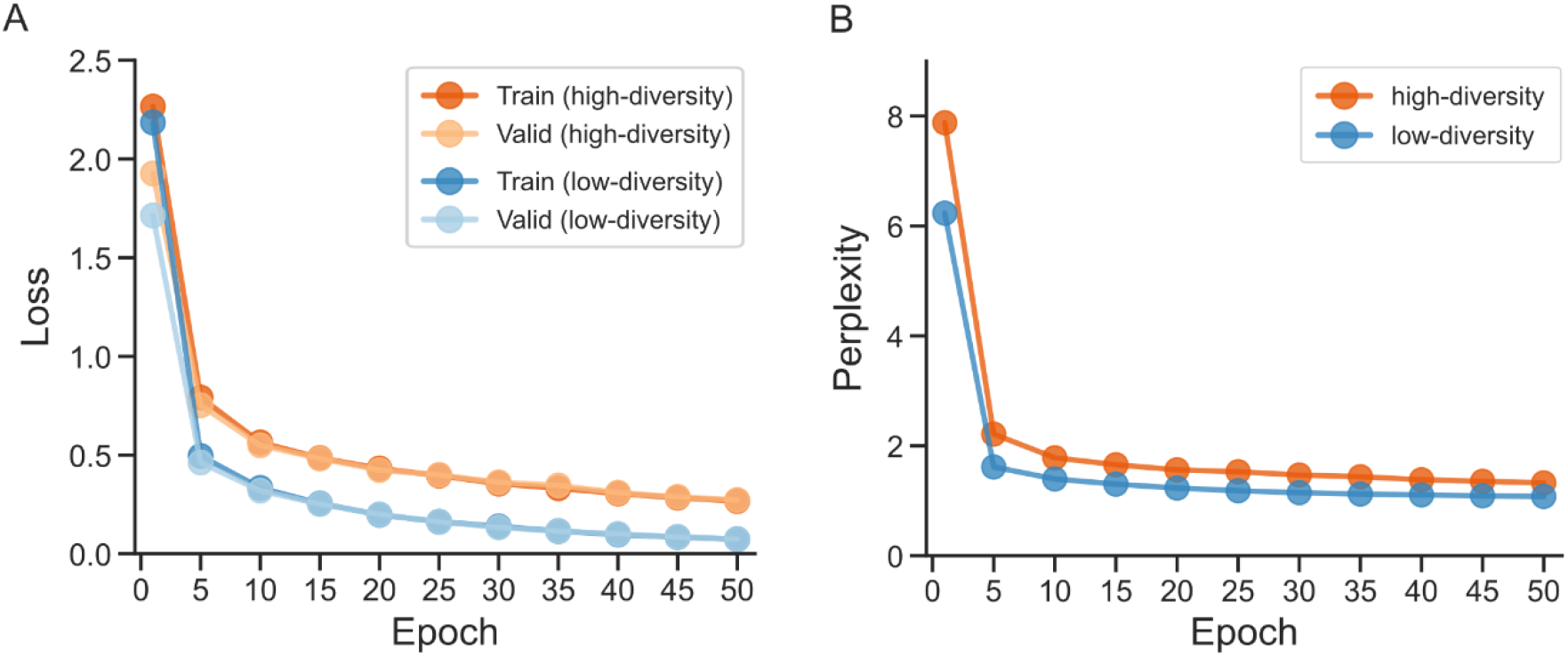
Learning curves for fine-tuning ESM2-35M. (A) Learning loss of training and validation data. (B) Perplexity of validation data.

**S3 Fig.**
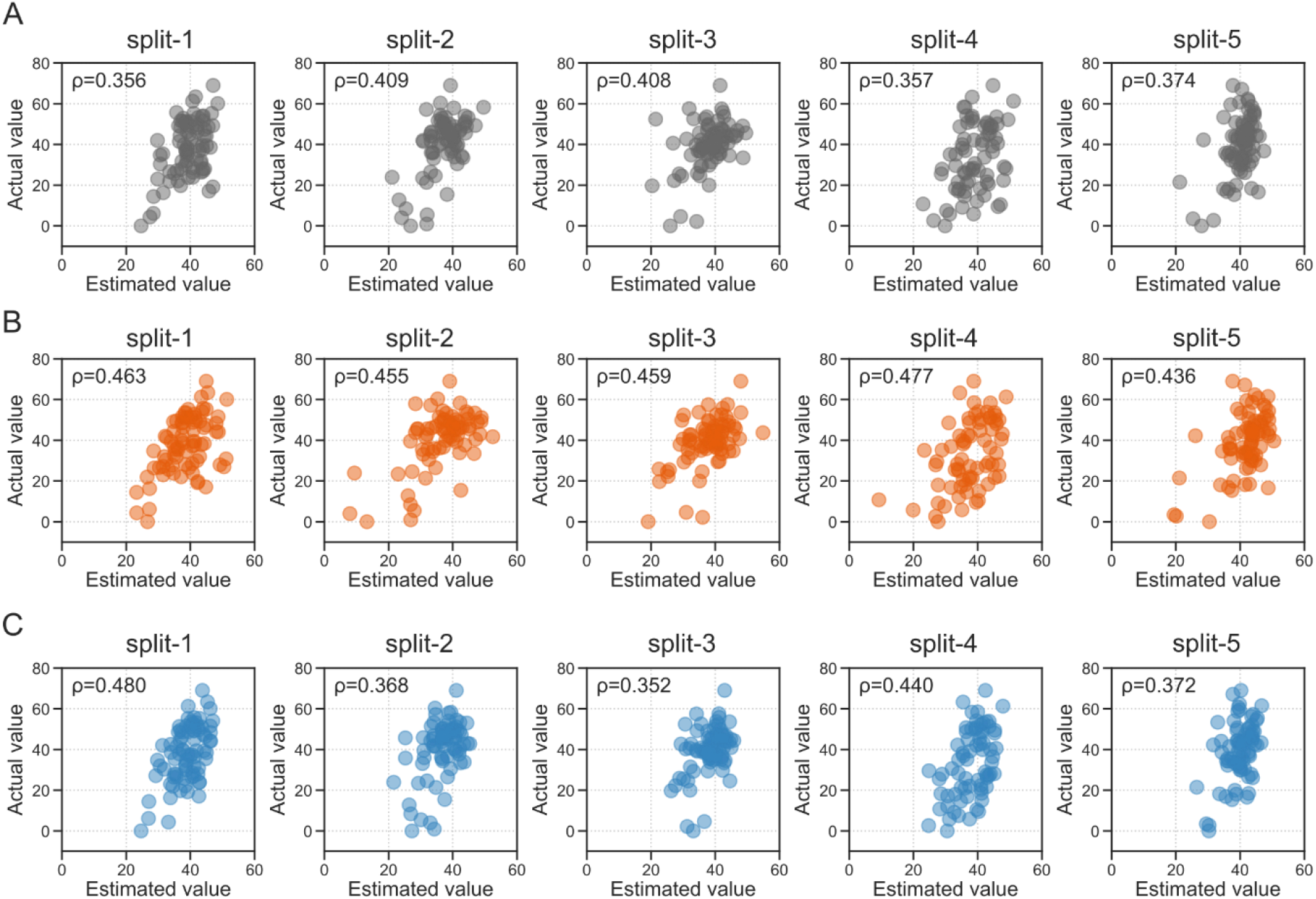
Comparison between estimated and actual values in prediction using pre-trained and fine-tuned ESM2-35M. (A-C) Plots show prediction results of validation data in 5-fold cross-validation (split-1~5). The plots represent pre-trained ESM-35M (gray), ESM2-35M_H/e10_ (orange), and ESM2-35M_L/e5_ (blue) from the top raw. The x-axis represents the estimated values, and the y-axis represents the actual values. The ρ value in the upper left on each plot indicates the Spearman’s rank correlation coefficient between the estimated and actual values.

**S4 Fig.**
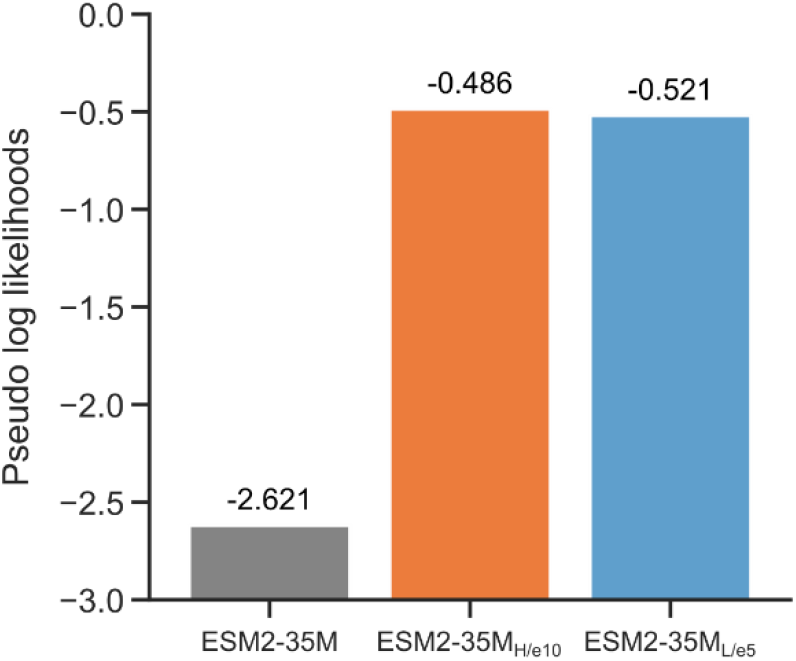
Pseudo log-likelihood of CAR mutants with pre-trained and fine-tuned ESM2-35M. The fine-tuned ESM2-35M with the highest prediction performance were used.

**S5 Fig.**
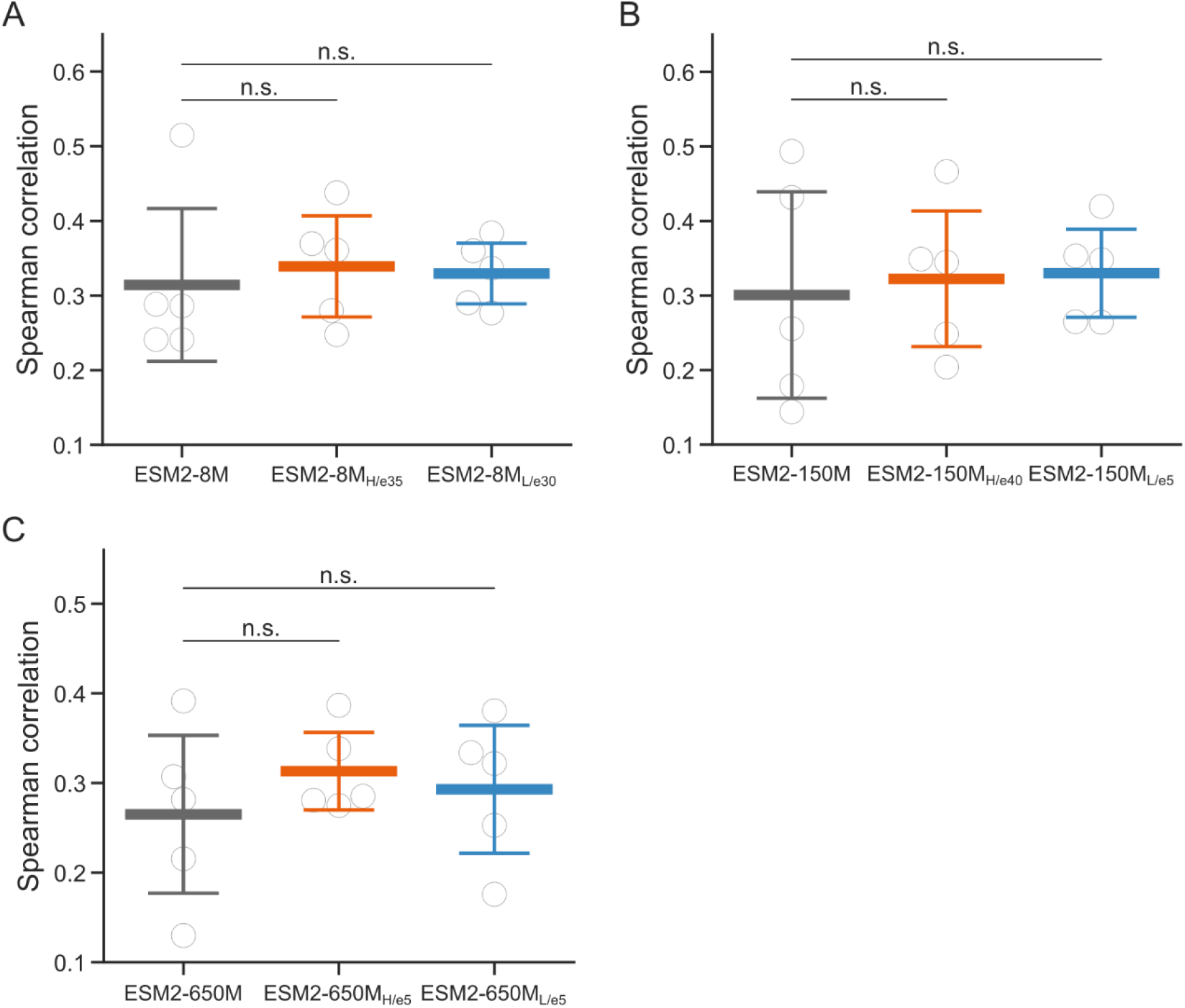
Comparison of the Spearman’s rank correlation coefficient between pre-trained and fine-tuned ESM2-8M, ESM2-150M, and ESM2-650M. (A-C) The circles represent individual Spearman’s rank correlation coefficient in each split of 5-fold cross-validation, and the lines indicated mean ± SD. The results of the fine-tuned ESM2-8M (A), ESM2-150M (B), and ESM2-650M (C) were taken at the maximum point within the range of 5 to 50 epochs. White circles represent individual values for each model of cross-validation. The thick middle lines represent the mean values, and the thin upper and lower lines indicate the mean ± SD. Multiple comparisons were performed using Dunnett’s test, n.s. (not significant).

**S6 Fig.**
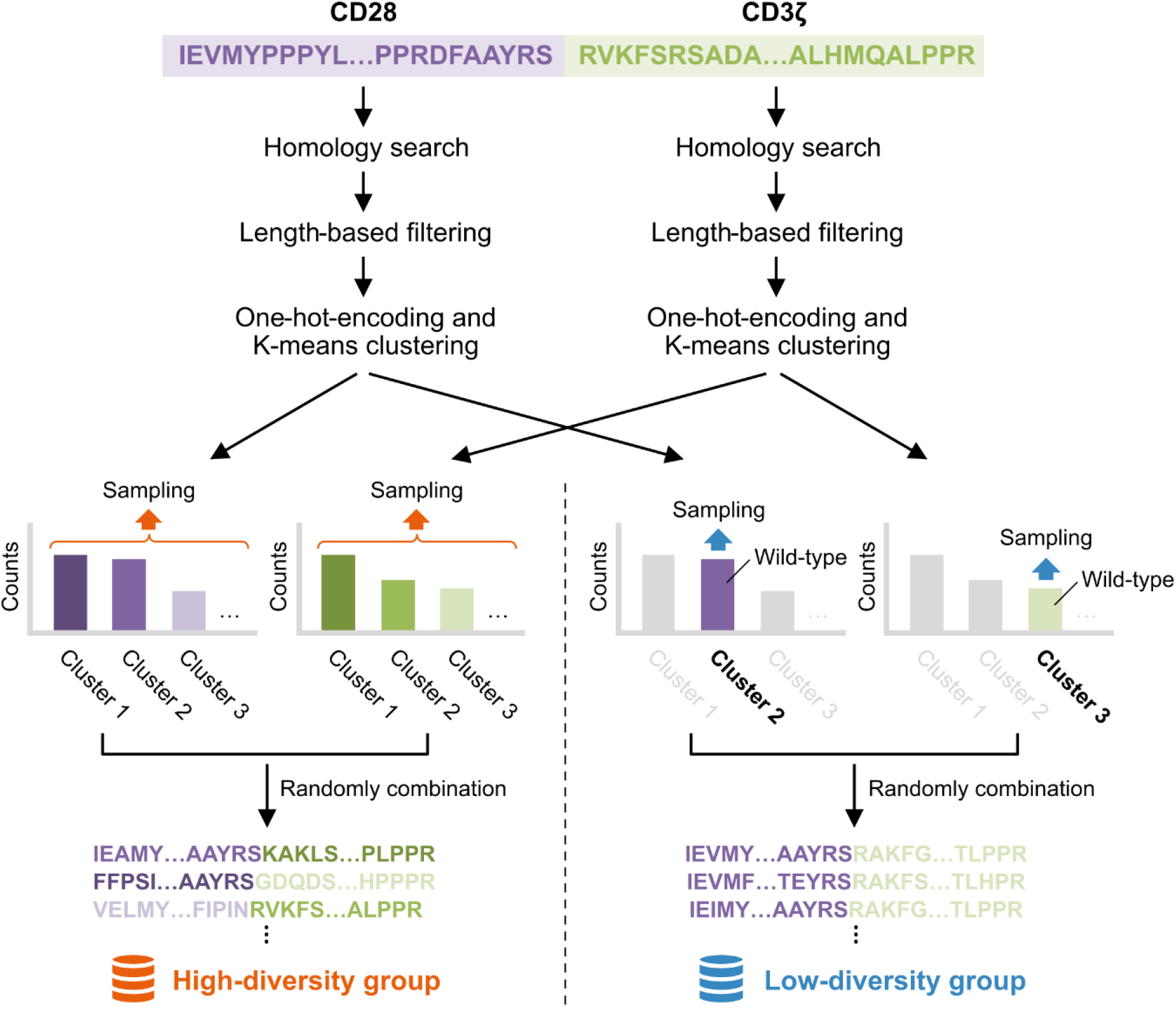
Sequence augmentation scheme. The target CAR regions in this study contain parts of CD28 and CD3ζ sequences. We created two types of groups with different diversity, using homology search and clustering methods (details provided in Materials & Methods).

**S1 Table.**
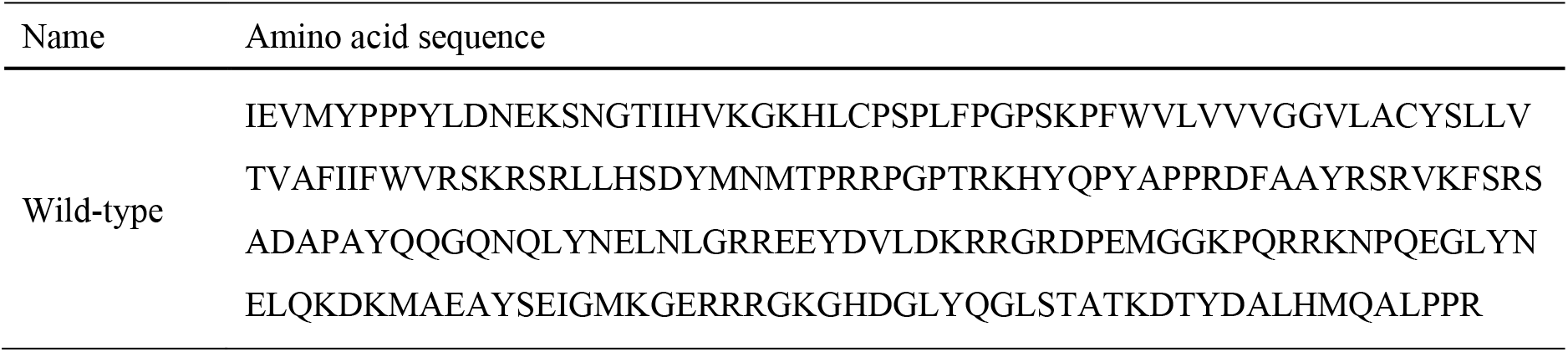
Amino acid sequence of wild-type.

**S2 Table.** Pre-trained ESM2 used in this study.

| Model name | Parameters | Layers | Hidden size | HuggingFace checkpoint name |
| --- | --- | --- | --- | --- |
| ESM2-8M | 8M | 6 | 320 | esm2_t6_8M_UR50D |
| ESM2-35M | 35M | 12 | 480 | esm2_t12_35M_UR50D |
| ESM2-150M | 150M | 30 | 640 | esm2_t30_150M_UR50D |
| ESM2-650M | 650M | 33 | 1280 | esm2_t33_650M_UR50D |

**S3 Table.** Summary of prediction performance using pre-trained and fine-tuned ESM2-8M, ESM2-150M, and ESM2-650M.

| Model | Epoch | Spearman $\rho$ | | Recall | | Precision | |
| --- | --- | --- | --- | --- | --- | --- | --- |
|  |  | Mean | SD | Top5 | Top10 | Top5 | Top10 |
| ESM2-8M | - | 0.314 | 0.114 | 0.083 | 0.124 | 0.320 | 0.240 |
| ESM2-8M<sub>H/e35</sub> | 35 | 0.339 | 0.076 | 0.094 | 0.165 | 0.360 | 0.320 |
| ESM2-8M<sub>L/e30</sub> | 30 | 0.329 | 0.045 | 0.073 | 0.134 | 0.280 | 0.260 |
| ESM2-150M | - | 0.301 | 0.155 | 0.104 | 0.209 | 0.400 | 0.400 |
| ESM2-150M<sub>H/e40</sub> | 40 | 0.322 | 0.102 | 0.083 | 0.125 | 0.320 | 0.240 |
| ESM2-150M<sub>L/e5</sub> | 5 | 0.330 | 0.066 | 0.083 | 0.229 | 0.320 | 0.440 |
| ESM2-650M | - | 0.265 | 0.098 | 0.073 | 0.177 | 0.280 | 0.340 |
| ESM2-650M<sub>H/e5</sub> | 5 | 0.313 | 0.048 | 0.104 | 0.209 | 0.400 | 0.400 |
| ESM2-650M<sub>L/e5</sub> | 5 | 0.293 | 0.080 | 0.082 | 0.218 | 0.320 | 0.420 |

## Footnotes

1 Since our focus is on constructing CAR-specific prediction model and not revealing the correlation between performance and model sizes, we leave comprehensive analysis for future work.

## References

[1] M. Sadelain, R. Brentjens, and I. Rivière, “The Basic Principles of Chimeric Antigen Receptor Design,” Cancer Discovery, vol. 3, no. 4, pp. 388–398, Apr. 2013, doi: 10.1158/2159-8290.CD-12-0548.

[2] S. T. Bhaskar, B. Dholaria, B. N. Savani, S. Sengsayadeth, and O. Oluwole, “Overview of approved CAR-T products and utility in clinical practice,” Clinical Hematology International, vol. 6, no. 4, pp. 100–106, Oct. 2024, doi: 10.46989/001c.124277.

[3] N. Bouchkouj, D. Przepiorka, and L. A. Fashoyin-Aje, “Obecabtagene Autoleucel for B-Cell Acute Lymphoblastic Leukemia,” JAMA, Feb. 2025, doi: 10.1001/jama.2024.28312.

[4] F. Korell, T. R. Berger, and M. V. Maus, “Understanding CAR T cell-tumor interactions: Paving the way for successful clinical outcomes,” Med, vol. 3, no. 8, pp. 538–564, Aug. 2022, doi: 10.1016/j.medj.2022.05.001.

[5] S. M. Albelda, “CAR T cell therapy for patients with solid tumours: key lessons to learn and unlearn,” Nat Rev Clin Oncol, vol. 21, no. 1, pp. 47–66, Jan. 2024, doi: 10.1038/s41571-023-00832-4.

[6] J. H. Park et al., “Long-Term Follow-up of CD19 CAR Therapy in Acute Lymphoblastic Leukemia,” New England Journal of Medicine, vol. 378, no. 5, pp. 449–459, Feb. 2018, doi: 10.1056/NEJMoa1709919.

[7] K. M. Cappell and J. N. Kochenderfer, “Long-term outcomes following CAR T cell therapy: what we know so far,” Nat Rev Clin Oncol, vol. 20, no. 6, pp. 359–371, Jun. 2023, doi: 10.1038/s41571-023-00754-1.

[8] A. Mei, “Engineering next-generation chimeric antigen receptor-T cells_ recent breakthroughs and remaining challenges in design and screening of novel chimeric antigen receptor variants,” Current Opinion in Biotechnology, 2024.

[9] R. B. Di Roberto et al., “A Functional Screening Strategy for Engineering Chimeric Antigen Receptors with Reduced On-Target, Off-Tumor Activation,” Molecular Therapy, vol. 28, no. 12, pp. 2564–2576, Dec. 2020, doi: 10.1016/j.ymthe.2020.08.003.

[10] P. Sharma et al., “Structure-guided engineering of the affinity and specificity of CARs against Tn-glycopeptides,” Proceedings of the National Academy of Sciences, vol. 117, no. 26, pp. 15148– 15159, Jun. 2020, doi: 10.1073/pnas.1920662117.

[11] J. C. Boucher et al., “CD28 Costimulatory Domain–Targeted Mutations Enhance Chimeric Antigen Receptor T-cell Function,” Cancer Immunology Research, vol. 9, no. 1, pp. 62–74, Jan. 2021, doi: 10.1158/2326-6066.CIR-20-0253.

[12] J. Feucht, “Calibration of CAR activation potential directs alternative T cell fates and therapeutic potency,” Nature Medicine, vol. 25, 2019.

[13] J. C. Greenhalgh, S. A. Fahlberg, B. F. Pfleger, and P. A. Romero, “Machine learning-guided acyl-ACP reductase engineering for improved in vivo fatty alcohol production,” Nat Commun, vol. 12, no. 1, p. 5825, Oct. 2021, doi: 10.1038/s41467-021-25831-w.

[14] Z. Wu, S. B. J. Kan, R. D. Lewis, B. J. Wittmann, and F. H. Arnold, “Machine learning-assisted directed protein evolution with combinatorial libraries,” Proceedings of the National Academy of Sciences, vol. 116, no. 18, pp. 8852–8858, Apr. 2019, doi: 10.1073/pnas.1901979116.

[15] Y. Saito et al., “Machine-Learning-Guided Library Design Cycle for Directed Evolution of Enzymes: The Effects of Training Data Composition on Sequence Space Exploration,” ACS Catal., vol. 11, no. 23, pp. 14615–14624, Dec. 2021, doi: 10.1021/acscatal.1c03753.

[16] K. K. Yang, Z. Wu, C. N. Bedbrook, and F. H. Arnold, “Learned protein embeddings for machine learning,” Bioinformatics, vol. 34, no. 15, pp. 2642–2648, Aug. 2018, doi: 10.1093/bioinformatics/bty178.

[17] A. S. Schwartz et al., “Deep Semantic Protein Representation for Annotation, Discovery, and Engineering,” Jul. 10, 2018, bioRxiv. doi: 10.1101/365965.

[18] E. C. Alley, G. Khimulya, S. Biswas, M. AlQuraishi, and G. M. Church, “Unified rational protein engineering with sequence-based deep representation learning,” Nat Methods, vol. 16, no. 12, pp. 1315–1322, Dec. 2019, doi: 10.1038/s41592-019-0598-1.

[19] A. Rives et al., “Biological structure and function emerge from scaling unsupervised learning to 250 million protein sequences,” Proceedings of the National Academy of Sciences, vol. 118, no. 15, p. e2016239118, Apr. 2021, doi: 10.1073/pnas.2016239118.

[20] A. Elnaggar et al., “ProtTrans: Toward Understanding the Language of Life Through Self-Supervised Learning,” IEEE Transactions on Pattern Analysis and Machine Intelligence, vol. 44, no. 10, pp. 7112–7127, Oct. 2022, doi: 10.1109/TPAMI.2021.3095381.

[21] N. Ferruz, S. Schmidt, and B. Höcker, “ProtGPT2 is a deep unsupervised language model for protein design,” Nat Commun, vol. 13, no. 1, p. 4348, Jul. 2022, doi: 10.1038/s41467-022-32007-7.

[22] D. Wang, F. Ye, and H. Zhou, “On Pre-trained Language Models for Antibody,” Mar. 01, 2023, arXiv: 2301.12112. doi: 10.48550/arXiv.2301.12112.

[23] S. Sledzieski, M. Kshirsagar, M. Baek, R. Dodhia, J. Lavista Ferres, and B. Berger, “Democratizing protein language models with parameter-efficient fine-tuning,” Proceedings of the National Academy of Sciences, vol. 121, no. 26, p. e2405840121, Jun. 2024, doi: 10.1073/pnas.2405840121.

[24] R. Schmirler, M. Heinzinger, and B. Rost, “Fine-tuning protein language models boosts predictions across diverse tasks,” Nat Commun, vol. 15, no. 1, p. 7407, Aug. 2024, doi: 10.1038/s41467-024-51844-2.

[25] C. Gordon, A. X. Lu, and P. Abbeel, “Protein Language Model Fitness Is a Matter of Preference,” Oct. 03, 2024, bioRxiv. doi: 10.1101/2024.10.03.616542.

[26] S. Biswas, “Low-N protein engineering with data-efficient deep learning,” Nature Methods, vol. 18, 2021.

[27] H. Yamaguchi and Y. Saito, “Evotuning protocols for Transformer-based variant effect prediction on multi-domain proteins,” Briefings in Bioinformatics, vol. 22, no. 6, p. bbab234, Nov. 2021, doi: 10.1093/bib/bbab234.

[28] S. Vincoff, S. Goel, K. Kholina, R. Pulugurta, P. Vure, and P. Chatterjee, “FusOn-pLM: a fusion oncoprotein-specific language model via adjusted rate masking,” Nat Commun, vol. 16, no. 1, p. 1436, Feb. 2025, doi: 10.1038/s41467-025-56745-6.

[29] Z. Lin et al., “Evolutionary-scale prediction of atomic-level protein structure with a language model,” Science, vol. 379, no. 6637, pp. 1123–1130, Mar. 2023, doi: 10.1126/science.ade2574.

[30] S. R. Eddy, “Accelerated Profile HMM Searches,” PLoS Comput Biol, vol. 7, no. 10, p. e1002195, Oct. 2011, doi: 10.1371/journal.pcbi.1002195.

[31] L.-C. S. Wang et al., “Targeting Fibroblast Activation Protein in Tumor Stroma with Chimeric Antigen Receptor T Cells Can Inhibit Tumor Growth and Augment Host Immunity without Severe Toxicity,” Cancer Immunology Research, 2014.

[32] O. Ueda et al., “Entire CD3ε, δ, and γ humanized mouse to evaluate human CD3–mediated therapeutics,” Sci Rep, vol. 7, no. 1, p. 45839, Apr. 2017, doi: 10.1038/srep45839.

[33] R. Zhang, J. Zhang, X. Zhou, A. Zhao, and C. Yu, “The establishment and application of CD3E humanized mice in immunotherapy,” Experimental Animals, vol. 71, no. 4, pp. 442–450, 2022, doi: 10.1538/expanim.22-0012.

[34] R. M.-H. Velasco Cárdenas et al., “Harnessing CD3 diversity to optimize CAR T cells,” Nat Immunol, vol. 24, no. 12, pp. 2135–2149, Dec. 2023, doi: 10.1038/s41590-023-01658-z.

[35] V. Levenshtein, “Binary codes capable of correcting deletions, insertions, and reversals,” Soviet physics. Doklady, 1965, Accessed: Feb. 28, 2025. [Online]. Available: https://www.semanticscholar.org/paper/Binary-codes-capable-of-correcting-deletions%2C-and-Levenshtein/b2f8876482c97e804bb50a5e2433881ae31d0cdd

[36] R. Gontijo-Lopes, S. J. Smullin, E. D. Cubuk, and E. Dyer, “Affinity and Diversity: Quantifying Mechanisms of Data Augmentation,” Jun. 04, 2020, arXiv: 2002.08973. doi: 10.48550/arXiv.2002.08973.

[37] F.-Z. Li, A. P. Amini, Y. Yue, K. K. Yang, and A. X. Lu, “Feature Reuse and Scaling: Understanding Transfer Learning with Protein Language Models,” Feb. 08, 2024, bioRxiv. doi: 10.1101/2024.02.05.578959.

[38] Z. Lin et al., “Language models of protein sequences at the scale of evolution enable accurate structure prediction,” Jul. 21, 2022, bioRxiv. doi: 10.1101/2022.07.20.500902.

[39] B. Chen et al., “xTrimoPGLM: Unified 100B-Scale Pre-trained Transformer for Deciphering the Language of Protein,” Dec. 09, 2024, arXiv: 2401.06199. doi: 10.48550/arXiv.2401.06199.

[40] Q. Fournier, R. M. Vernon, A. van der Sloot, B. Schulz, S. Chandar, and C. J. Langmead, “Protein Language Models: Is Scaling Necessary?,” Sep. 23, 2024, bioRxiv. doi: 10.1101/2024.09.23.614603.

[41] K. Janocha, A. Ling, A. Godson, Y. Lampi, S. Bornschein, and N. Y. Hammerla, “Harnessing Preference Optimisation in Protein LMs for Hit Maturation in Cell Therapy,” Dec. 03, 2024, arXiv: 2412.01388. doi: 10.48550/arXiv.2412.01388.

[42] A. Paszke et al., “PyTorch: An Imperative Style, High-Performance Deep Learning Library,” in Advances in Neural Information Processing Systems, Curran Associates, Inc., 2019. Accessed: Feb. 28, 2025. [Online]. Available: https://papers.nips.cc/paper_files/paper/2019/hash/bdbca288fee7f92f2bfa9f7012727740-Abstract.html

[43] T. Wolf et al., “Transformers: State-of-the-Art Natural Language Processing,” in Proceedings of the 2020 Conference on Empirical Methods in Natural Language Processing: System Demonstrations, Q. Liu and D. Schlangen, Eds., Online: Association for Computational Linguistics, Oct. 2020, pp. 38–45. doi: 10.18653/v1/2020.emnlp-demos.6.

[44] A. Wang and K. Cho, “BERT has a Mouth, and It Must Speak: BERT as a Markov Random Field Language Model,” Apr. 09, 2019, arXiv: 1902.04094. doi: 10.48550/arXiv.1902.04094.

